# Distributed Phase Oscillatory Excitation Efficiently Produces Attractors Using Spike Timing Dependent Plasticity

**DOI:** 10.1101/2020.10.22.351379

**Authors:** Eric C. Wong

## Abstract

The brain is thought to represent information in the form of activity in distributed groups of neurons known as attractors, but it is not clear how attractors are formed or used in processing. We show here that in a randomly connected network of simulated spiking neurons, periodic stimulation of neurons with distributed phase offsets, along with standard spike timing dependent plasticity (STDP), efficiently creates distributed attractors. These attractors may have a consistent ordered firing pattern, or become disordered, depending on the conditions. We also show that when two such attractors are stimulated in sequence, the same STDP mechanism can create a directed association between them, forming the basis of an associative network. We find that for an STDP time constant of 20ms, the dependence of the efficiency of attractor creation on the driving frequency has a broad peak centered around 8Hz. Upon restimulation, the attractors self-oscillate, but with an oscillation frequency that is higher than the driving frequency, ranging from 10-100Hz.

## INTRODUCTION

The brain is generally thought to represent abstractions, such as objects or events, in the form of groups of active neurons, referred to variously as cell assemblies or ensembles [1–5], engrams [6, 7], cliques [8], or attractors [9]. These groups (or subgroups thereof) are consistently active when the brain is engaged in processing related to the corresponding abstraction. Networks composed of attractors have been proposed as models of learning, memory, and cognition [3, 10–13]. While there is evidence for the existence of attractors in the brain [7], the mechanisms by which they are formed, utilized, and connected to one another remain unclear.

The spike timing dependent plasticity (STDP) mechanism [14, 15] is well established, and increases synaptic weight when a postsynaptic neuron fires within a short time window after the firing of a presynaptic neuron, while it weakens the synaptic weight when the firing order is reversed. With controlled excitation of a sequence of neurons in a network, this mechanism can form nearly arbitrary patterns of directed connections. Note that if a series of neurons is excited with a time spacing that is conducive to STDP, in a circular pattern such that the first neuron is re-excited after the last, a circular set of directed connections can be created by STDP. This can be the basis for the construction of circular attractors. When excitation forms a repeating sequence, the excitation incident to each neuron will be periodic, but must also have a full range of phase offsets across the neurons of the attractor.

We hypothesize that distributed phase periodic excitation may be a general mechanism used by the brain to construct attractors.

Oscillatory activity at a broad range of frequencies has been observed in the brain and has been linked to many functions including learning and plasticity [16, 17], processing across brain areas [18, 19], and place encoding [20, 21]. Sensitivity to the phase of these oscillations is also well established, particularly in the theta [20, 22–24] and gamma [18, 25, 26] bands, and theta and gamma oscillations are thought to be coordinated with one another through phase relationships [12, 27–32]. The hippocampus in particular is known to be a source of phase modulated theta oscillations [33].

Attractors in the form of a linear linked list, known as synfire chains have been studied as potential functional structures in the brain [34], and have been shown to demonstrate compositionality [35]. Circular attractors [36, 37] and synfire chains [38] have been shown to emerge slowly but spontaneously with random inputs while under the influence of STDP.

We focus here on the potential use of structured inputs by the brain to construct and associate circular attractors. We simulate spiking neural networks with a simplified but biologically plausible architecture and parameters, and report the following key findings:

1. Distributed phase oscillating excitation of an excitatory network with inhibitory feedback and standard STDP rapidly produces circular attractors that can be re-excited.
2. For an STDP time constant of 20ms, the dependence of attractor formation efficiency on driving frequency displays a broad peak centered around 8Hz.
3. The natural oscillation frequency of the resulting attractors is higher than the driving frequency during formation, and typically in the range of 10-100Hz
4. The resulting attractors may be either circular or disordered.
5. The resulting attractors can be associatively linked to one another after formation using the same STDP mechanism as that which constructed them.

## METHODS

Computer simulations were performed using the spiking neural simulator Brian 2 [39], and all code used to produce the data and figures for this article are available at (https://github.com/spindragon/thetagamma). Our simulated network consisted of 10,000 excitatory neurons (E), 1,600 inhibitory neurons (I) with E-E, E-I, and I-E synaptic connections and was simulated with a 0.01ms time step. In concept the E and I neurons corresponded to pyramidal and basket cells, respectively. Default parameters for the networks are summarized in **Table 1**.

**Table 1:** Model Parameters

| Parameter | Symbol | Value |
| --- | --- | --- |
| Membrane time constant | $\tau_{mem}$ | 10ms |
| Adaptation time constant | $\tau_{adapt}$ | 100ms |
| Refractory period | $\tau_{refract}$ | E: $6.3 \pm 1.7ms$<br>I: $5 \pm 1ms$ |
| STDP time constant | $\tau_{STDP}^{+(-)}$ | 20ms |
| Resting potential | $V_{rest}$ | -70mV |
| Threshold potential | $V_{thresh}$ | -50mV |
| Adaptation potential increment | $\Delta V_{adapt}$ | E: 6mV<br>I: 5mV |
| Reset potential | $V_{reset}$ | -65mV |
| Gaba potential | $V_{gaba}$ | -80mV |
| Peak connection probability | $\rho_0$ | EE: 0.8<br>EI: 0.4<br>IE: 0.3 |
| Lorentzian radius<br>(in units of grid spacing) | $\lambda$ | EE: 30<br>EI: 2<br>IE: 5 |

Leaky integrate and fire neurons with adaptation were simulated with dynamics defined by:

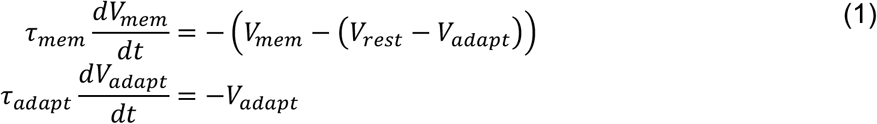

Where *V_mem_* and *τ_mem_* are the membrane potential and time constant, *V_rest_* is the resting membrane potential, *V_adapt_* is a dynamic variable that implements a spike driven adaptation with time constant *τ_adapt_*. The membrane potential decays toward the quantity (*v_rest_ − V_adapt_*), while *V_adapt_* is incremented by spikes and decays toward zero with time constant *τ_adapt_*. The membrane potential was not allowed to decrease below the reversal potential of GABA, implementing the idea that inhibitory input decreases the membrane potential toward this potential.

Neurons spiked when *V_mem_* reached a threshold potential of *V_threshold_*, and at this time:

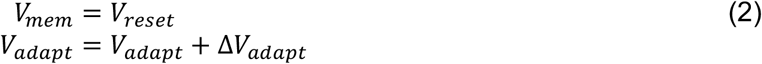

The membrane potential remained at *V_reset_* for a refractory period *τ_refract_* which was taken from Gaussian distributions with mean±std of 6.3 ± 1.7*ms* [40] for E and 5 ± 1*ms* for I.

A conduction delay between E neurons was set to the sum of an axonal component that was calculated from the Euclidean distance between neurons with an assumed conduction velocity of 5m/s, and a dendritic component that was randomly taken from a uniform distribution between 2ms and 4ms [41]. Conduction delays between E and I neurons were set to 0. When the spike potential reaches the target neuron, the membrane potential of the postsynaptic neuron is incremented by:

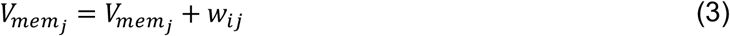

where *w_ij_* is the synaptic weight between neurons *i* and *j*. With this formulation, synaptic weights are in units of mV, and were capped at *w_max_* = 5*mV*. Since there was a 20mV difference between resting and threshold potentials, approximately 4 or more simultaneous incoming spikes are required in order to trigger a postsynaptic spike.

The connections between E neurons were subject to STDP and implemented learning, while E-I and I-E synapses had fixed synaptic weights that were chosen empirically to provide stability to the network (values in Table 1). When STDP was active and a postsynaptic spike occured in neuron *j* at time Δ*t* relative to the arrival of the presynaptic potential from neuron *i*. a candidate change Δ*w_ij_* in the synaptic weight was created and given by:

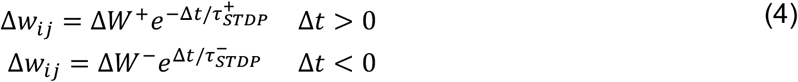

Where Δ*W*^+(−)^ are maximum possible weight changes and 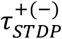 are the time constants for potentiation and (depression). Various values of Δ*W*^+^ were tested, but the ratio of depression to potentiation was set to Δ*W*^−^/Δ*W*^+^ = −1.05, following [15]. A ratio slightly larger in magnitude than one (-1.1) was also found empirically in [42], and implements a gradual decay of synaptic weights in the presence of random inputs. A ratio equal to or greater than 1.0 is necessary to prevent divergence of weights. The time constants for STDP were set to 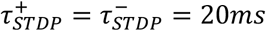. Unless otherwise noted, these candidate weight changes were accumulated during the training period and applied prior to the test period, following the observation that STDP appears to take effect in the seconds to minutes following training, rather than immediately [43, 44].

Neurons were placed on a square 2D grid with 1cm or 15cm sidelength to approximately simulate mouse and human brain dimensions, respectively. In order to avoid edge effects the network topology was toroidal, meaning that opposite edges are connected. The connection probabilities *ρ* were defined by a Lorentz function of the Euclidean distance *r* between neurons, given by:

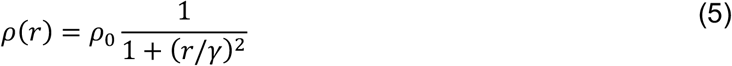

Where *ρ*_0_ is the probability at *r* = 0 and *γ* is the radius at which the probability is reduced by a factor of 2. The Lorentzian is a common long-tailed function and was chosen to emphasize local connections, but with sparse longer range connections as well. Connections were more spatially distributed for E-E connections, and more localized for E-I and I-E connections, with parameters in Table 1. No self-connections were allowed in the E network.

An attractor was defined as a group of *N_a_* = 500 neurons randomly selected from the 10,000 neuron E network, each labeled with sequential neuron number *n* upon selection. During training, these neurons were subject to a Poisson distributed spiking input with a peak rate of 200Hz, where each input spike increases the membrane potential by 50mV, sufficient to excite any neuron that is not refractory. For a given driving frequency *f* (period 1/*f*), each neuron experienced a half cosine windowed spiking rate *C*(*t*) for a fraction *d* (duty cycle) of each period. The timing of excitation was shifted linearly from 0 to 1/f across the neurons of the attractor, generating a phase rotating or circular excitation. Finally, the excitation function was multiplied by a window function Ω(*t*) to mitigate any spurious effects of a sudden onset or cessation of stimulation. The full excitation function *X* is given by:

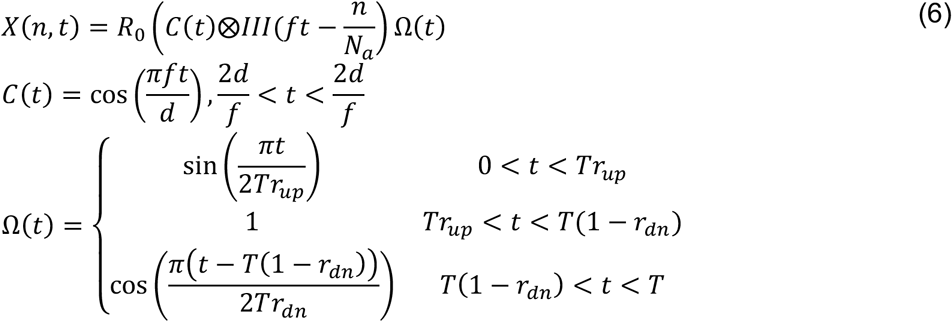

where *R*_0_ = 200*Hz* is the peak spiking rate, *III*(*t*) is the comb or shah function, *T* is the total duration of excitation, and *r_up_* = 0.1 and *r_dn_* = 0.1 are the fractions of *T* occupied by sinusoidal ramps up and down from the plateau, respectively. Poisson sampling from this rate function was taken at 1ms intervals. An example of the spikes produced by the excitation function *X*(*n, t*) is shown in **Figure 1a**. Note that because of the random selection of neurons comprising the attractor, there is no functional difference between sequential excitation of the attractor neurons (time offset linearly related to neuron number *n* as in equation 6 above) and excitation with random but uniformly distributed time offsets. There is also no spatial order to the attractor.

**Figure 1:**
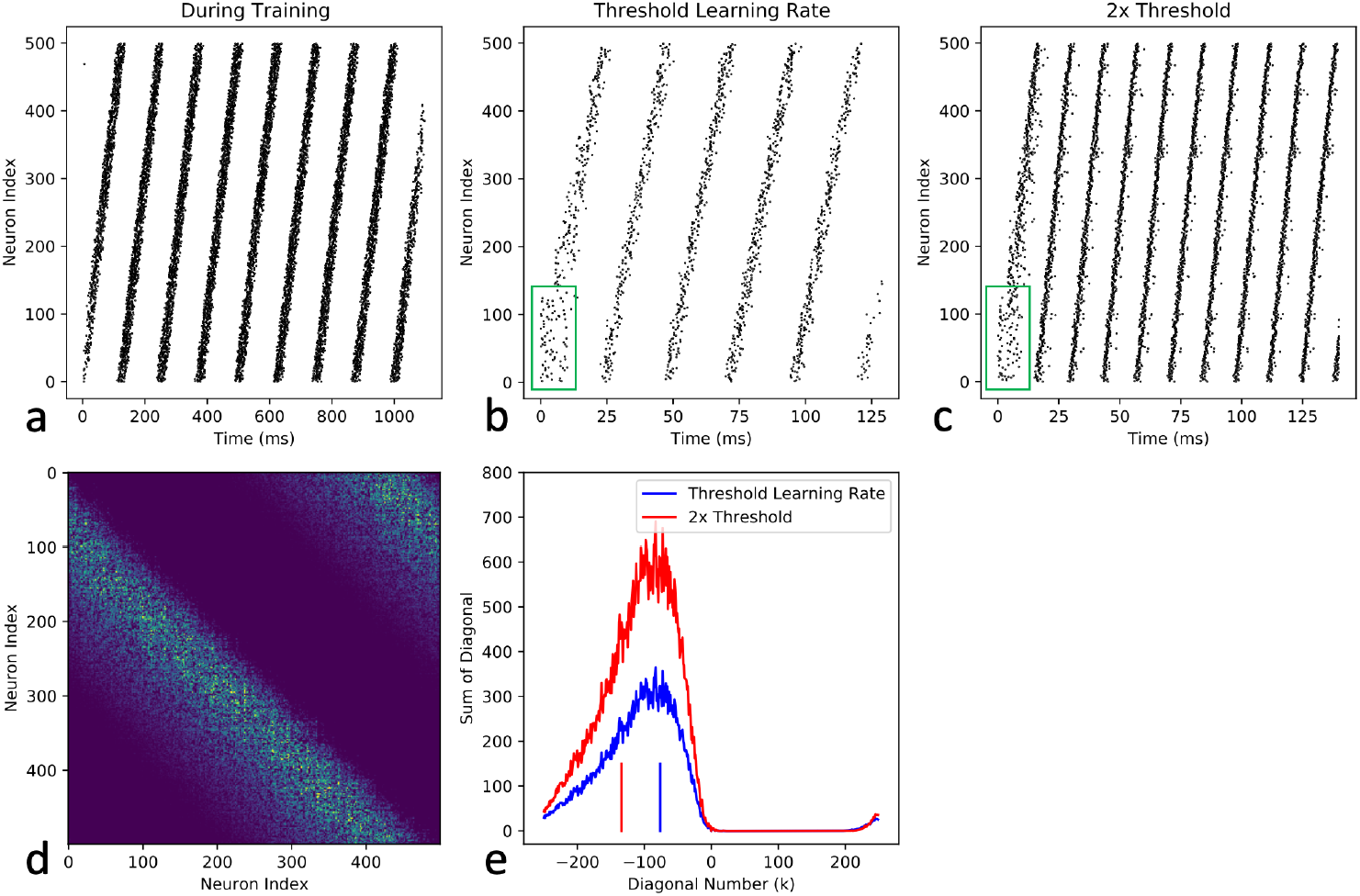
Distributed phase oscillating excitation constructs oscillating attractors. a) Spikes in attractor neurons during 8Hz rotating phase stimulation. b) Response to re-excitation of first 25% of attractor neurons for 10ms (green box) after learning at the minimum learning rate that results in self-oscillation upon re-stimulation. Self-oscillation frequency is 42Hz. c) Response to re-excitation after learning at twice the threshold learning rate. Self-oscillation frequency is 74Hz. d) Learned weight matrix for E-E synapses between attractor neurons after training, showing a prominent band of increased weights below the diagonal, indicating that neurons have learned to excite other neurons within the attractor that are ahead of them in index number. e) Projection of diagonals of weight matrix learned at threshold and twice threshold learning rates, accounting for wraparound due to circular attractors (see text). The approximate location of the peak at threshold learning rate can be calculated from the observed oscillation frequency and the conduction delay (see text), and is indicated by the blue vertical line. Higher oscillation frequency at twice threshold learning rate corresponds to weights at a more negative diagonal (red vertical line), which corresponds approximately to the diagonal at which the weights at 2x threshold are equal in magnitude to those at the peak of the weights at threshold (see text).

For estimation of the self-oscillation frequency, attractors were re-excited using 10ms of stimulation of the first 25% of neurons in the attractor and the frequency was estimated for the period from 20-120ms after the beginning of re-excitation. The spike record over this time period was divided into 40 time bins of Δ*t* =2.5ms each, and for each bin a complex representation of the activity *A*(*t*) was calculated as:

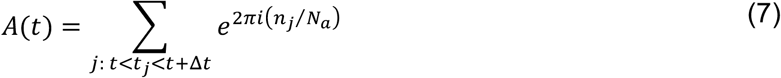

where *n_j_* and *t_j_* are the neuron index and time for spike *j*. *A*(*t*) represents the status of the attractor as a point in the complex plane, with one rotation in this plane corresponding to a cycle of the attractor, and |*A*| provides a metric of the relative SNR of the local phase estimate. The frequency was calculated as the slope of a linear least squares fit to the unwrapped phase of *A*(*t*), weighted by |*A*|.

## RESULTS

After training, attractors can generally be re-excited using a brief period of excitation of a subset of the neurons in the attractor. As an example, we trained a 1cm network using 8Hz circular excitation at 25% duty cycle for 1.1s, with a learning rate given by Δ*W*^+^= 0.128*mV* using the stimulus shown in **Figure 1a**. The candidate weight changes were applied after training, and the attractor was re-excited by exciting the first 25% of the nodes in the original training sequence for 10ms. After re-excitation, the attractor self-oscillated at 42Hz, in approximately the same order as during training (**Figure 1b**). The learning rate above was the lowest that would support self oscillation for at least 120ms upon re-excitation. We refer to this as the threshold learning rate 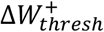 for future reference, but note that this rate is strongly dependent upon the details of the excitation pattern, and is not a fundamental physiological quantity. **Figure 1c** shows the response upon re-excitation for 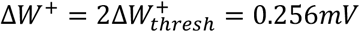, demonstrating a higher self-oscillation frequency of 74Hz. The weight matrix learned at 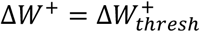 is shown in **Figure 1d** for synapses between neurons within the attractor, with indices sorted in the same order as the training excitation (vertical axis of **Figures 1 a-c**). Note that there is a prominent band of increased weights below the diagonal, indicating that each neuron has learned to excite neurons with a higher index number. The increased weights in the upper right corner reflect the circular nature of the attractor, indicating that neurons near the end of the attractor excite those near the beginning.

**Figure 1e** shows the sum of each diagonal of the weight matrix for offsets *k* where *N_a_*/2 < *k* < *N_a_*/2, with the weight matrix assumed to be replicated to the left and to the right (vertical replication is equivalent). Using this construction each point represents the sum of all weights connecting any presynaptic neuron to any neuron -k neurons ahead of it in the circular attractor. The position of the band of increased weights is related to the self-oscillation frequency of the attractor. For each neuron, there is a conduction delay *τ_d_*, that represents the transmission time from dendritic synapse to axonal synapse. If the neuron participates in transmission around a circular attractor, then the product of *τ_d_* and the oscillation frequency gives the fraction *ϕ* of the circle spanned by transmission through a neuron:

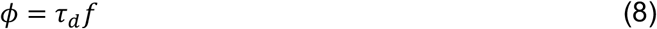

In the example above the mean value of *τ_d_* was 3.6ms, giving *ϕ* = (3.6*ms*)(42*Hz*) = 0.15. If neurons in the attractor are exciting neurons that are ahead in phase by *ϕ*, there should be large synaptic weights at diagonal

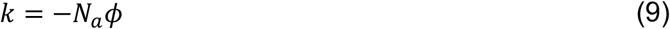

since −*k/N_a_* also represents the fractional rotation around the attractor. The blue vertical line in the Figure marks this diagonal (k=−76), and is located near the peak of the learned weights at the threshold learning rate. This calculation is a rough estimate of the relationship between the weight matrix and the oscillation frequency as it uses the mean conduction delay as a first order description of the distribution of delays, but provides a reasonably consistent estimate nevertheless. For the higher learning rate (red trace in **Figure 1e**) the measured self-oscillation frequency was significantly higher (74Hz), and we interpret this as follows. At a higher learning rate the weights simply scale linearly relative to lower learning rates until the weights begin to clip at *w_max_*, thus the red curve in **Figure 1e** is scaled but not shifted along *k* relative to the blue curve. At the threshold learning rate, the weights at the peak of the blue curve are presumably just sufficient to excite the group of neurons −*k_peak_* neurons ahead in the attractor. If this is the case, then for higher weights, there is a broad range of values of *k* for which the weights are sufficient to propagate excitation. However, when the neurons at the farthest end of the excitable neurons (most negative *k*) are excited, those neurons will continue to propagate the wave of excitation, and the self-oscillation will proceed at the highest supportable frequency. We note that the higher observed oscillation frequency of 74Hz corresponds to *k* = −133 (vertical red line in the figure) using Equations 8 and 9 above, and aligns approximately with the point where the higher learning rate weight matches the peak of the weights at threshold.

In order to characterize the dependence of the self oscillation frequency on the driving frequency, we repeated the above simulation for driving frequencies from 1-20Hz, network sizes of 1cm and 15cm, and stimulation duty cycles *d* = [0.125,0.25,0.5]. Results are shown in **Figure 2**. The self-oscillation frequency was positively correlated with the driving frequency, but the dependence was not simply proportional. For the 1cm network, the self-oscillation frequency was found to exceed 100Hz for the highest driving frequencies, while the 15cm network was limited by conduction delays to approximately 40Hz. For each parameter set we found the threshold learning rate 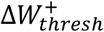, and measured the oscillation frequency at threshold and twice threshold, as above (**Figure 2a**). We also define a learning efficiency *η*:

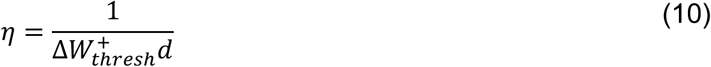

where the denominator is proportional to the product of the change in synaptic weight per appropriately timed spike pair times the number of input spikes used in that simulation, and thus to the sum total of synaptic weight required to construct an attractor. The efficiency *η* reports the inverse of this sum, and is shown in **Figure 2b** normalized to the maximum value within the entire data set. Note that there is a broad peak in *η* for driving frequencies between approximately 3Hz and 15Hz, centered around 8Hz, while the self-oscillation frequencies (**Figure 2a**) are also broadly distributed but largely occupying the gamma band. The profiles of the learned weight matrices for the two network sizes and the two extremes of driving frequencies are shown in **Figure 2c** for d=0.25 and the threshold learning rate. Using a mean conduction delay of *τ_d_* = 3.6*ms* for the 1cm network and 12.5*ms* for the 15cm network, and the observed self-oscillation frequencies, the expected peak weight offsets *k* were estimated using Equations 8 and 9 above and shown as vertical bars in the figure.

**Figure 2:**
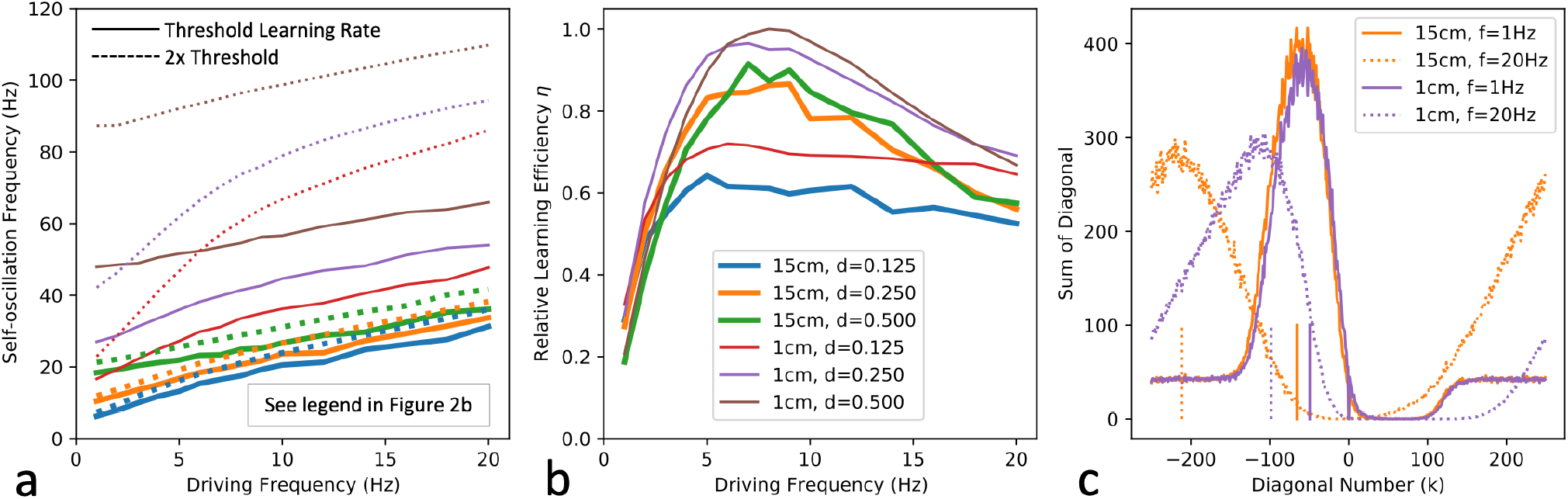
Frequency dependence of attractor construction and self-oscillation. a) Self-oscillation frequency vs driving frequency during training. (see **Figure 2b** for color legend). The observed self-oscillation frequency is uniformly higher than the training frequency, and increases with a higher learning rate (dotted lines). b) The relative learning efficiency *η* is defined as the inverse of the total weight change required to maintain self-oscillation (see text). For both small and large networks there is a broad peak in efficiency centered around a driving frequency of 8Hz. c) Diagonals of learned weight matrices are shown for the extremes of driving frequency. Expected location of weight peaks, calculated using Eqs. 8 and 9 and the observed self-oscillation frequencies are shown, and are consistent with the observed weights.

In the above examples, the activity within the attractor after training oscillates with approximately the same circular pattern as the driving oscillator, but at a higher frequency. However, under some circumstances the self-sustained activity in the attractor becomes disordered rather than periodic, and two examples are shown in **Figure 3**. In the first example, we start with the 15cm network trained with a driving frequency of 20Hz, a duty cycle of 0.25, and twice the threshold learning rate. The learned weight changes were applied before re-excitation but no additional weight changes were applied. After re-excitation the network oscillates for a short period of time, but within 200ms, there is a transition to a firing pattern that is no longer periodic (**Figure 3a-b**). The final 100ms of the 1s period is shown in detail, demonstrating an apparently chaotic pattern of firing. Thus we see that even when STDP is only active during controlled periodic stimulation, the resulting attractor can become disordered. The learned weight matrix is shown in **Figure 3c**, and the profile of the diagonals of this matrix is also the dotted orange trace in **Figure 2c**. The broad peak of the weight matrix is nearly *N_a_*/2 removed from the diagonal, suggesting that the excitation may be bouncing back and forth across the circular attractor rather than traveling in a well-defined circular wave. In a second example we start with a 1cm network trained with a driving frequency of 8Hz and a duty cycle of 0.25. The learning rate was set high, at 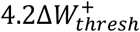, and the STDP weight changes were applied immediately, rather than accumulated and applied after the training epoch. Stimulus was applied for 500ms, and the network continued to fire and evolve after the stimulus was stopped (**Figure 3d-e**). Around 250ms into the stimulation the effects of plasticity are apparent as the driven neurons begin to fire other neurons with a frequency that is higher than the driving frequency. Before the end of the stimulation at 500ms, the learned synapses have taken over the firing of the attractor. The final 100ms of the epoch is shown in detail in **Figure 3e** and shows a similar chaotic pattern of firing as in the previous example. However, in this case the learned weight matrix (**Figure 3f**) is void of any obvious diagonally banded structure.

**Figure 3:**
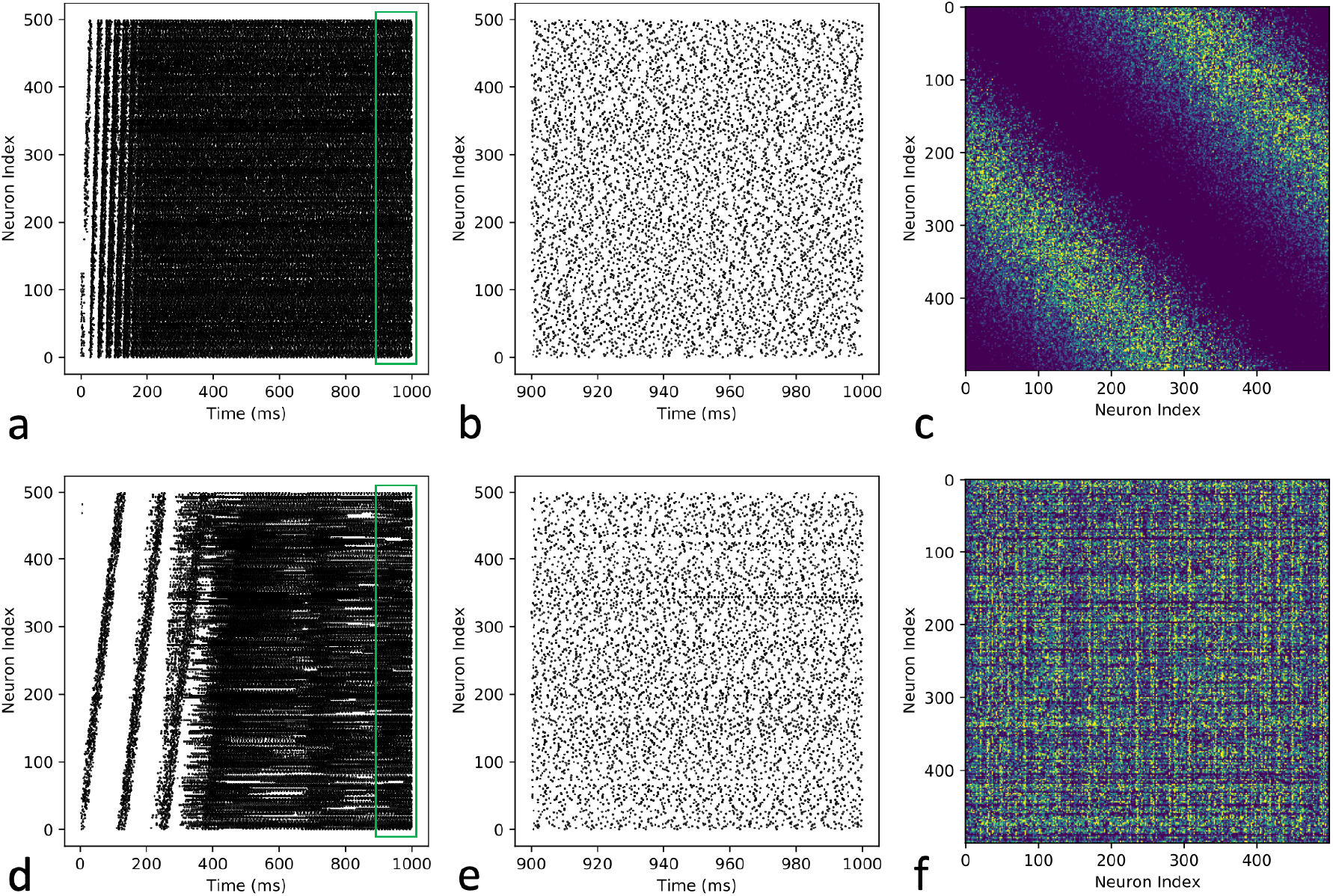
Disordered attractors. a) Circular attractor trained with 20Hz driving frequency, with weight changes applied after training and unchanged thereafter. Training period is not shown. The attractor was re-excited and allowed to evolve for 1s, demonstrating initial oscillation, evolving to a disordered state. b) Final 100ms (green box in a) in detail, showing non-circular and non-periodic firing. c) Weight matrix shows bands of high weights near *k = N_a_*/2, such that neurons excite neurons nearly opposite themselves in the circular attractor. d) Attractor with weight changes applied immediately both during training and ongoing thereafter. 8Hz stimulation was applied for 500ms, but after approximately 250ms, learned weights begin to dominate the firing. e) Final 100ms of 1s epoch in detail, showing a similar pattern of non-circular and non-periodic firing. f) Learned weight matrix is devoid of obvious diagonal banding, though we do not know whether a permutation of indices would restore a banded structure.

In addition to forming new attractors, we asked whether the same STDP mechanism can associate attractors with one another [1]. We separately trained three attractors in a 1cm network using the same method as in Figures 1 and 2, with a driving frequency of 8Hz, a learning rate Δ*W*^+^= 0.16*mV*, which is 1.4 times the threshold learning rate. Neurons for each attractor were independently and randomly chosen with overlap allowed. Spikes during this training are shown in **Figure 4a**. The apparently spurious spikes in the attractors not being trained represent neurons that are present in two or more attractors. After training, the learned weight changes were applied, and each attractor was re-excited in sequence 140ms apart, with the resulting spike train shown in Figure 4b. Note that each attractor is excitable while the previous attractor is still oscillating, and extinguishes the activity of the previous attractor. During this epoch, STDP was active with a learning rate given by Δ*W*^+^= 0.04*mV*, accumulating candidate weight changes which were applied after this second epoch. In a third epoch, only the first attractor was excited, and excites the next two in sequence, demonstrating directed associations between the attractors. A similar example is shown for disordered attractors in **Figures 4d-f**. In this example each of three attractors in a 1cm network were trained at 8Hz and 0.25 duty cycle for 500ms using a high learning rate Δ*W*^+^= 0.48*mV*, with immediate application of STDP changes, resulting in immediate generation of disordered attractors. For this example, non-overlapping attractors were used because a 5% overlap as in the example above was found to cause immediate excitation of all attractors when one was excited, effectively merging the attractors. Each attractor was allowed to evolve for an additional 600ms after stimulation under ongoing STDP to strengthen the synaptic weights within the attractors. Between the three training periods in this epoch, the network was reset to a resting state before the next training period. As in the previous example, during a second training epoch the three attractors were then re-excited in sequence, 140ms apart, but with STDP ongoing at a learning rate of Δ*W*^+^=0.024*mV* and immediately applied (**Figure 4e**). In a third epoch, the first attractor alone was re-excited, and led to the rapid sequential excitation of the next two attractors (**Figure 4f**). These findings of sequential linking of attractors are reproducible, but not robust to large changes in parameter settings in our simulations. Under many circumstances attractors will fail to excite one another or newly excited attractors will fail to extinguish previous attractors, but we show these examples to demonstrate that directional associations between attractors can be formed using the same STDP mechanism as that which formed the attractors.

**Figure 4:**
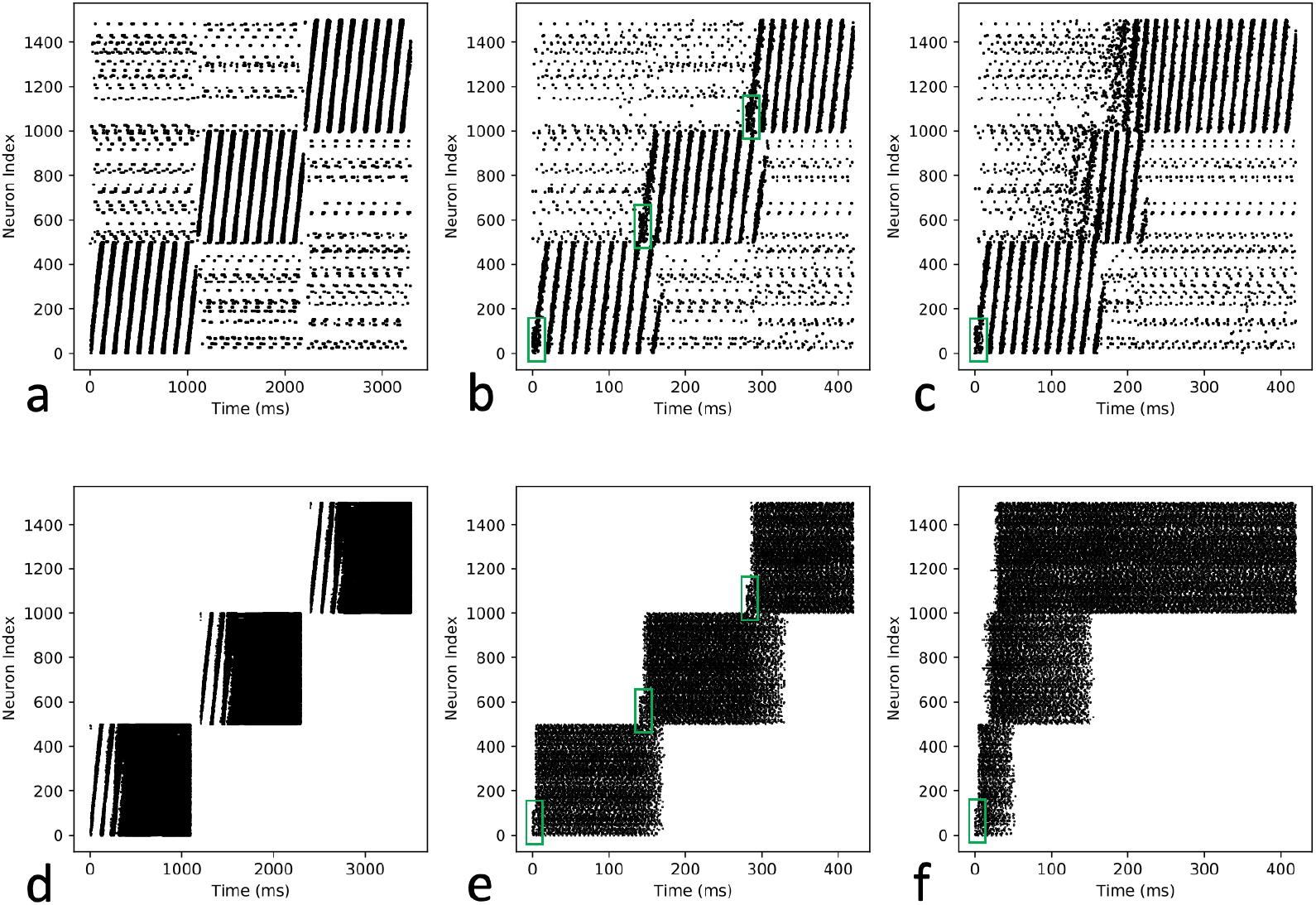
Linking of attractors. a) Three overlapping attractors were sequentially trained. Off diagonal spikes represent neurons that are in more than one attractor (5% overlap). Weight changes were applied after training. b) Each attractor was re-excited in sequence, 140ms apart, with STDP active, and weight changes were applied after training. Green boxes indicate spikes associated with reexcitation. Note that the excitation of the second and third attractors extinguish the activity of the previous attractors. c) Only the first attractor was then re-excited, and had learned to subsequently excite the second and third attractors. d) Three non-overlapping attractors were trained with STDP applied immediately, under the same conditions as in **Figure 3d-f**. Activity of the network was halted at the end of each 1.1s training epoch. e) Each attractor was re-excited in sequence, 140ms apart, with STDP active and applied immediately. Again, the excitation of the second and third attractors extinguish the activity of the previous attractors. f) Only the first attractor was re-excited, and quickly excites the second and third attractors in sequence, with variable time required to extinguish the previous attractors.

## DISCUSSION

### Theta band excitation creates gamma band attractors

We have shown that in a realistic simulated spiking network, excitation of random neurons using oscillating excitation with distributed phase offsets efficiently construct circular attractors using STDP, that high efficiency occurs with driving frequencies across a broad range centered around the theta band, and that the resulting attractors oscillate in the gamma band. We use the term efficiency to describe the net change in synaptic weights for a given number of input spikes. We believe that the construction of attractors described here is efficient because the timing of excitation is well matched to the temporal characteristics of the STDP process. With phase modulated oscillatory input, any pair of neurons in the target network that fire close together in time do so consistently in the same order. Therefore, because of the approximate antisymmetry of the STDP response curve, each synapse between such pairs will experience only potentiation or only depression of the synaptic weight, and plasticity mechanisms will make modifications quickly. Any process that stimulates pairs of neurons with inconsistent order will produce a mixture of potentiation and depression, partially cancelling the effects, and therefore requiring more STDP driven events per unit net change in weights. Stimulation using oscillations without distributed phase, or using random schedules of excitation does not produce consistent asymmetry in the firing order between any pair of neurons, and for antisymmetric STDP result in much slower net modification of synaptic weights, although such stimulation has also been shown to form attractors [37, 38].

We believe that the increase in self-oscillation frequency relative to the driving frequency during training is due to the fact that the STDP time constant is relevant at training, but not directly relevant during self-oscillation. During training the sum of the STDP time window and the conduction delay, along with the driving frequency, define the potential range of phase rotations *ϕ* for connected neuron pairs. However, during self-oscillation the same learned phase rotation per neuron pair will be effective, but will be traversed in a time given by only the conduction delay without an additional STDP related delay [45], resulting in a higher oscillation frequency. Consistent with this idea, we have verified that increasing the STDP time constant results in higher self-oscillation frequencies (data not shown). Note that the positive correlation between STDP time constant and the self-oscillation frequency is atypical for the relationship between time constants and frequencies within most systems, which are more commonly inversely related, or negatively correlated.

### Attractors may be circular or disordered

The attractors formed can remain circular when self-oscillating after training if STDP is not active or not consolidated by subsequent neuromodulatory activity. However, it appears that ongoing STDP generally causes the attractors to evolve to a disordered state [46]. We have not determined whether these disordered attractors would demonstrate a circular structure with an appropriate permutation of the indices for the neurons in the attractor. Searching for such a permutation is a combinatorial problem that we have not yet addressed, and similar problems are known to be NP hard [47]. We do note that even within a stable circular attractor the firing of each node is not precisely periodic [48], and the attractor is better described by a wave of excitation travelling around the attractor but with disordered excitation within the wave, much like a wave of applause that can travel around a stadium consistently and periodically without perfect periodicity from any one person. Whether the brain utilizes attractors in circular vs disordered configurations would depend on the degree to which STDP or other plasticity mechanisms are available and consolidated as the attractor is re-stimulated and used for processing. However, we also note that the distinction between circular and disordered attractors may not be important functionally, as it appears that re-excitation and association of attractors does not seem to depend critically on the pattern of activity (see **Figure 4**).

### Associative links between attractors can be created using STDP

In our simulations, attractors can be associated with one another using the same STDP mechanism as that which constructed the attractor in a single training example, such that the excitation of one attractor leads to excitation of the next, analogous to the ‘phase sequence’ described by Hebb [1]. These associations are directional, and we hypothesize that the increased frequency of self-oscillation may be instrumental in allowing for these directional associations to form. For the formation of attractors, driving frequencies in the theta range allow for the construction of directed connections around the circular attractor because the driving oscillation is slow enough to allow for several STDP time constants to elapse during a circumnavigation of the attractor, and therefore for several sequential subsets of neurons to be connected in series. After attractor formation, frequency acceleration into the gamma range allows for one or more oscillations *within one* STDP time constant, which enables many (or all) neurons of one attractor to connect by STDP to many neurons of the next attractor in a single rapid transition from one to the next. With slower oscillation, only a subset of one attractor would be connected to the next in a rapid transition. If attractors are associated during a period of overlap that is long compared to the STDP time constant, then the association would naturally become bidirectional.

### Limitations of this work

The simulations shown here are simplified and neglect many details that may be important for the quantitative analysis of the effects reported. For example, the numbers of neurons and synapses are unrealistically low, and were chosen for computational speed. The typical number of synapses on the dendritic tree of a pyramidal cell is approximately 25,000 [49], vs approximately 3400 in our simulations. Because of the low number of synapses, we set the maximum synaptic weight to 5*mV* so that approximately 5 incoming spikes can trigger a post-synaptic spike, whereas the threshold number of incoming spikes in vivo is likely closer to 135 [49]. Small network size also increases the overlap between attractors that are randomly chosen, and we found in the example in **Figure 4d-f** that the 5% overlap of randomly selected attractors was sufficient to cause the attractors to effectively merge into one. Dendritic computation is also likely highly complex [50], but was reduced to simple integration in our model. Our feedback inhibition system was composed of static E-I and I-E connections and weights, while in vivo there are many cell types that likely implement many homeostatic and dynamically modulated feedback systems as well as pruning mechanisms to facilitate learning and maintain stability under a wide range of conditions. Despite these simplifications, we tried to use physiologically realistic numbers for the parameters that we believed would have the largest effect on the frequency dependence of training and self-oscillation, such as the STDP time constant, conduction delays, and refractory periods.

### Summary

We propose here that phase modulated theta oscillations may be used by the brain to rapidly construct attractors, and that the effective range of driving frequencies is closely tied to the STDP time constant through the phase rotation per neuron around the attractor. We show that the acceleration of the self-oscillating frequency of the resulting attractors into the gamma range is both a natural consequence of the learning mechanism and fortuitous for the subsequent association of attractors. We note that simply broadcasting a phase modulated theta oscillation through a random set of connections to any network that has latent synapses and STDP active can produce a distributed attractor. For example, this may be a key component of consolidation of memories from the hippocampus, which is known to generate phase modulated theta oscillations [22, 33, 51], to a sparse distributed representation in the cortex. Note that such an attractor generated in isolation would have no meaning since it would simply be a random set of neurons bound together. However, when excited in coordination with other attractors, a newly formed attractor could develop associations with those other attractors by the mechanisms proposed here, and thereby learn to take on the same meaning as the originating hippocampal source.

## ACKNOWLEDGEMENTS

We would like to thank Richard Buxton, Thomas Liu, and Peter Bandettini for many useful discussions regarding this work, and for helpful review of this manuscript.

## Notes

### Competing Interest Statement

The authors have declared no competing interest.

